# Spatial coding of conspecifics in subpopulations of hindbrain pyramidal cells of the gymnotiform electrosensory system

**DOI:** 10.1101/2023.08.09.552616

**Authors:** Oak E Milam, Gary Marsat

## Abstract

Localizing the source of a signal requires sophisticated neural mechanisms and we are still uncovering the coding principles that support accurate spatial processing. Weakly electric fish can detect and localize distant conspecifics, but the way this spatial information is encoded is unclear. Here, we investigate the spatial representation of conspecific signals in the hindbrain to determine how the properties of the heterogenous population of pyramidal cells affect the spatial coding accuracy of conspecific signals. We hypothesize that specific subsets of cells provide more accurate spatial information about conspecific location. We stimulated the fish with an artificial signal that replicates both the spatial and temporal structure of conspecific signals. We recorded from cells with various receptive field positions covering the entire body surface and analyzed the spike train with spike-train distance metrics to determine how accurately the location of the stimulus is encoded. We found that some pyramidal cells (such as ON-type, and those within the deep layer) encode the spatial information more accurately while other subgroups (OFF-type, and superficial layer) provide less accurate information. Our results help us understand how the heterogeneity of a population of cells allow the efficient processing of signals and suggest that a segregation of the spatial information stream starts earlier in the sensory pathway.

## Introduction

Nervous systems must accurately encode sensory information about environmental stimuli and a central goal of neuroscience is to reveal how this is accomplished efficiently (Barlow, 1961; Bialek and Rieke, 1992; Bullock et al., 2005; Dayan and Abbott, 2001; Shannon, 1953). Courting a signaling mate or surviving an agonistic encounter between a competitor are a couple of behavioral examples where encoding spatial information reliably is essential for piloting social interaction (Bradbury and Vehrencamp, 2011; Pedraja et al., 2016). Yet, how this spatial information is represented by populations of neurons to guide such behaviors remains poorly understood. Here, we investigate the spatial coding accuracy of heterogeneous pyramidal cell populations in the hindbrain collected via *in-vivo* electrophysiological recordings.

*Apteronotus leptorhynchus*, a gymnotiform wave-type weakly electric fish, produces a continuous, quasi-sinusoidal, electric organ discharge (EOD) via an electric organ located in the tail (Lissmann, 1951; Lissmann, 1958). The EOD drives the baseline discharge of tuberous electroreceptors (P-units) distributed across the entire body. P-units are the electroreceptor type most relevant for encoding electrosensory input, used for communicating with conspecifics and navigating the environment (Bullock, 1969; Bullock, 1982). The afferents from each P-unit provide trifurcated, unilateral input to different subtypes of pyramidal cells located in the maps of the electrosensory lateral line lobe (ELL): the lateral segment (LS), centro-lateral segment (CLS), and centro-medial segment (CMS; for review see Krahe and Maler, 2014; Milam et al., 2019). Multiple topographic maps in the ELL are comprised of a heterogeneous network of ON and OFF-type pyramidal cells (Heiligenberg and Dye, 1982; Maler, 2009a; Shumway et al., 1989). Pyramidal cells are organized in a columnar layout, containing superficial, intermediate, and deep-type pyramidal cells. Within each map, pyramidal cells vary in their response properties and center-surround receptive field parameters (Chacron et al., 2001; Krahe et al., 2008). Receptive fields in the LS map are the largest, the CMS map contains the smallest receptive fields, and receptive fields in the CLS map are intermediate. It has been suggested that different neural maps are specialized for certain behavioral tasks (Allen and Marsat, 2018; Maler, 2007). However, besides a few focused studies, little is known about how pyramidal cells (at the individual neuron or population level) respond to spatially realistic, conspecific stimuli (Kelly et al., 2008; Litwin-Kumar et al., 2012).

Recent field and lab studies on interacting weakly electric fish indicate that these animals possess an aptitude for detecting and localizing conspecific signals in their environment, even in conditions where sensory cues are limited (Zupanc and Maler, 1993; Stamper et al., 2012; Henninger et al., 2018; Knudsen, 1975; Berman and Maler, 1999; Yu et al., 2012; Fotowat et al., 2013; Jung et al., 2016). The diffuse nature of these signals (affecting peripheral receptors covering the entire body, i.e., “global signal”) suggests that the central nervous system must discriminate between small differences in the spatial signal to encode conspecific location accurately. Though behavioral observations clearly demonstrate their sensory capacity, how the nervous system accomplishes this task remains unknown. Our goal in this study is to understand how the primary electrosensory area of the nervous system encodes the location of a conspecific based on their self-generated signal. We aim to uncover how multiple receptor inputs are integrated by pyramidal cells in the hindbrain so that relevant spatial information is encoded in a way that enables accurate localization of conspecifics. We argue that efficient extraction of spatial information should involve implementing different neural streams and codes. We hypothesize that a heterogeneous population of pyramidal cells uses different neural coding strategies for efficiently processing conspecific stimuli that vary based on their spatial parameters.

In this study, we directly compare neural heterogeneity at the lowest level of the electric fish’s central nervous system – the ELL – and accurate neural coding of conspecific location. We first show that the spatially realistic, conspecific stimulus elicits responses from ELL pyramidal cells represented in both rate and temporal aspects of the spike train. After single cell analysis, we quantify a population response from a heterogeneous pool of pyramidal cells and demonstrate that information about the spatial stimulus is encoded in the pattern of the population response. We calculate the discrimination performance of the population and find that spatial stimuli are accurately and efficiently encoded. This confirms that combining sensory input from multiple pyramidal cell receptive fields yields higher discrimination performance for coding conspecific position. We separate the population into categories and reveal a pyramidal cell type specialization for spatial coding of conspecific information based on the aspect of the response used for discrimination. Finally, we assess neural coding performance across a range of stimulus spatial scales, and using a weighted analysis, confirm that a segregation of conspecific spatiotemporal information begins in the primary sensory area of weakly electric fish.

## Materials and Methods

### Animals

Wild-caught *Apteronotus leptorhynchus* were obtained from commercial fish suppliers.

Tanks water conductivity were maintained at 200-300 µS and at temperatures of 26-27°C. All procedures were approved by the West Virginia University IACUC.

### Electrophysiology

Surgical techniques were as previously described (Allen and Marsat, 2019; Marsat and Maler, 2010; Marsat et al., 2009). Briefly, *A. leptorhynchus* was anesthetized with tricane methanesulfonate (Western Chemical, Inc.) and respirated during surgery. A local anesthetic (Lidocaine HCL 2%, Hospira, Inc.) was applied, and the skin overlying the craniotomy site was removed. A fixed post with a circular opening was glued to a portion of the exposed skull for stability. The fish was immobilized with an injection of tubocurarine chloride pentahydrate (0.2 mg ml^-1^, TCl). The experimental tank contained water with conductivity at 250 (±10) µS and temperature at 26 (±1) °C. The portion of the skull above the ELL was removed. A cone was secured to the fixed post, allowing top-down access to the exposed ELL. Melted resin was used to form a watertight seal between the ventral opening of the cone and the skull around the exposed ELL. ACSF was applied to the brain. This cone allows for full body submersion into the experimental tank during recordings while preventing the brain from coming in contact with tank water. Respiration was switched from general anesthesia to anesthetic-free water for respiration. *In vivo*, single-unit recordings of the lateral segment (LS) and centrolateral segment (CLS) were performed using metal-filled extracellular electrodes (Frank and Becker, 1964). Recordings were amplified (A-M Systems, Model 1700) and data recorded (Axon Digidata 1500 and Axoscope software, Molecular Devices) at a 20kHz sampling rate. Pyramidal cells of the LS and CLS were identified based on the blood vessel landmarks, depth of penetration (in the dorsal-ventral plane), and response properties of the (Maler et al., 1991; Saunders and Bastian, 1984).

### Stimulation

All stimuli were sampled at 20 kHz and created in MATLAB (MathWorks, Inc.). Our stimulation procedure replicates the amplitude modulations (AM) experienced during social interactions. The baseline EOD was recorded between the head and tail of the fish. Each EOD cycle triggered a sine wave generator (Rigol DG1022A) to output one cycle of a sinusoidal signal with matching frequency to the fish’s EOD. This signal was then multiplied using a custom-built signal multiplier by the AM stimulus to create the desired modulation of the electric field. Stimuli were played through a custom made stimulus isolator into the experimental tank using one of three configurations: a global stimulation, via two 30.5 cm carbon electrodes arranged parallel to the longitudinal axis of the fish; a local stimulation, via two silver chloridized electrodes 0.5 cm apart positioned at various positions near the skin surface; an artificial conspecific stimulation (i.e., fishpole), via two silver chloridized electrodes embedded in agarose with a conductivity of 35 µS (Hupé and Lewis, 2008; Kelly et al., 2008; Walz et al., 2013). Stimulation for global and local configurations were adjusted to provide ∼20% contrast relative to the baseline EOD strength.

The fishpole signal was calibrated to match the amplitude of the electric field potential of real fish recorded from 24 different positions and distances around the fish (n=7, of mixed gender and size). One of the silver wires represented the rostral end of the fishpole (5.5 cm long), and the other silver wire represented the caudal end (0.1 cm long). The two wires were separated by 4 cm, yielding a fishpole with 9.6 cm in length and a zero-plane potential located at ∼70% of the rostral to caudal body length, in accordance with previous observations (Assad and Bower, 1997). The solidified, agarose body was carved to match the body shape of a real fish. The fishpole was positioned in the experimental tank at three different orientations/azimuths (0, 45, 90°), and 7 different locations around the fish 10 cm away. The 7 locations were the operculum, mid-body, and the zero-potential plane for both ipsilateral and contralateral sides of the fish (6 locations), and 1 location directly caudal to the fish (see Figure 1A). We use the term, orthogonal, to describe stimulus positions where the rostral end of the fishpole is oriented toward the receiving fish. Sinusoidal AM (SAM) stimuli were 40 s long, modulated at 30 Hz, and were played through the fishpole during the experiments.

**Figure 1.**
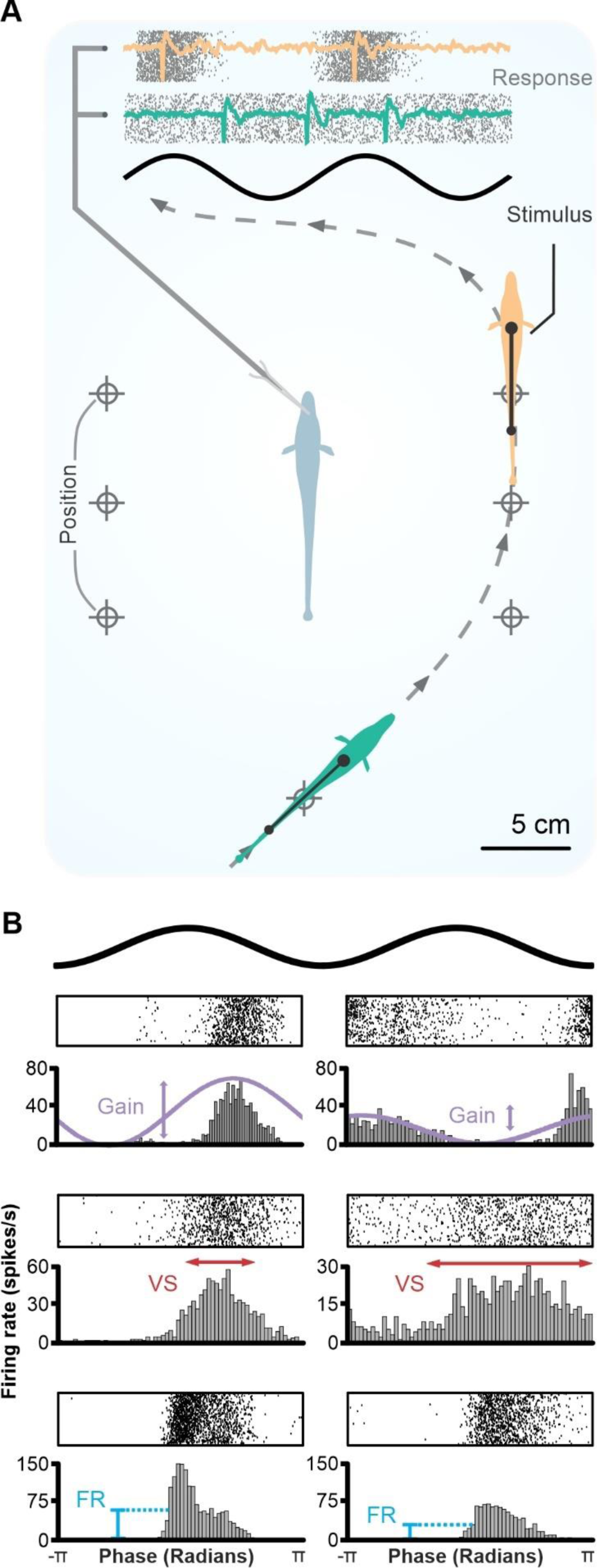
Spatially realistic conspecific signals and response strengths in the ELL. **(A)** Schematic of the experimental design. An immobilized *A. leptorhynchus* (center, light blue) is stimulated using a conspecific dipole mimic (i.e., “fishpole”) positioned at various spatial locations and azimuths around the experimental tank (at distances of 10 cm), while recording extracellularly from ELL pyramidal neurons *in vivo*. Neural responses to two cycles of the conspecific stimulus (top, black) are shown as raster plots layered with a trace of the raw neural recordings. The examples highlight differences in the pattern of the spike train responses for encoding spatial stimuli (orange and green). **(B)** Measurement of response strength to stimulations with conspecific stimulus from different stimulation sites (yellow site for A.: left column; green site: right column). Raster plots of the response are shown above, and the corresponding peristimulus time histogram (PSTH) are shown below. Changes in spatial location and orientation of the conspecific stimulus can elicit increases or decreases in the spike train response of ELL pyramidal cells. Responses to the spatial stimulus vary in stimulus-response gain (top, purple), vector strength (middle, red), and mean firing rate (bottom, blue).

### Data Analysis

All analyses described here were performed using MATLAB. Spike trains collected from experimental recordings were first binarized into a sequence of zeros (no spike) and ones (spike). The binarized sequence was transformed into instantaneous firing rates by convolution with a gaussian filter. We used either the binarized spike train or the instantaneous firing rate (see below) that were separated into 1 second, 50% overlapping segments. Statistical analyses were performed using the MATLAB statistical analysis toolbox and custom-made scripts. Data was tested using a 2-way (or n-way) ANOVA. Following all ANOVAs, post hoc comparisons were made using a Tukey-Kramer test. Mean differences were considered statistically significant when p < 0.05.

### Gain

The stimulus-response gain (*G*) to SAM stimulation was calculated by:

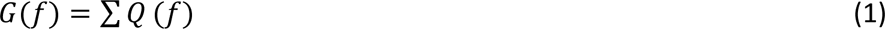

where *Q* is the power spectral density of the convolved spike train, *f* are the frequencies within ± 0.5 Hz of the target SAM frequency (30 Hz). A larger stimulus-response gain value indicates a larger response from the neuron to a stimulus at the target frequency.

### Vector Strength

The strength of phase locking to SAM stimulation was calculated by:

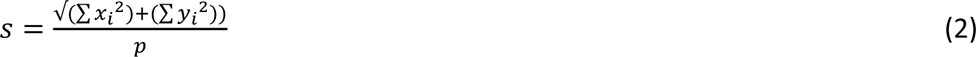

where *p* is the number of spikes, and *x* and *y* are the sine and cosine phases of the stimulus at which the *i* spike occurs (Goldberg and Brown, 1969; Marsat and Pollack, 2004). The vector strength, *s*, quantifies the precision and clustering of responses to a given phase of the stimulus cycle, with 0 being equal response at all phases of the beat, and 1 being a perfectly precise response at a single phase of the beat.

### Discrimination Analysis

Our discrimination analysis is based on a weighted Euclidean distance analysis that relates directly to the information carried by a population of neurons to discriminate between stimuli (see Marsat et al., 2023, for more details). Here, we compare stimuli from different locations and used one of three measurements to quantify response strength: mean firing rate, vector strength, or gain for each 1 s response segment. A weight is assigned to each neuron for each pair of stimuli being compared. The weight is based on the Kullback-Leibler divergence in the response distributions for each stimulus. The weight is normalized to 1 across neurons within a population response, the response strength is then multiplied by this weight. Population responses are then taken as data point in Euclidean space where each dimension is the weighted response of one neuron in the population. An ensemble of population response is thus considered, each of which is composed of a random 1 s segment of response from a subset of *n* neurons from the population (*n* will be varied, see below). The Euclidean distance between responses to the same stimulus and across different stimuli are then compared. Larger distances indicate less similarity between spike trains. Stimuli that can be easily discriminated will elicit responses that are very different (i.e., large Euclidean distance) relative to the variability across responses to the same stimulus. The weighting procedure allows to optimize the decoding efficiency by assigning a stronger contribution to the Euclidean distance to neurons that carry more information about the difference in the stimuli. The distributions of Euclidean distances for responses to the same stimulus *P(D_xx_)* and across the two stimuli being compared *P(D_xy_)* are then used in a receiver operating characteristic (ROC) analysis. Receiver operating characteristic (ROC) curves were generated by varying a threshold distance value T; for each threshold, the probability of non-discrimination (*P_D_*) is calculated as the sum of *P(D_xy_>T)* and the probability of false discrimination *(P_F_)* is calculated as the sum *P(D_xx_>T)*. The error probability is taken as the minimum error, *E,* across thresholds:

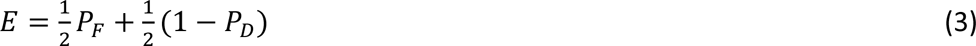

Error probability of 0.5 indicates chance-level discrimination while an error rate of 0 indicates that the responses are different enough to support perfectly accurate discrimination.

### Efficiency Rate

The size of the population of neurons used in the discrimination analysis can be varied. If it is based on the information contained in a single neuron, discrimination will be less accurate than if the information from many neurons is considered. By plotting the error probability as a function of the number of neurons included in the analysis, we can estimate how quickly the error rate decreases with increasing population size. This rate of decrease is representative of efficiency in population coding since it reflects how much information each neuron contributes. Based on this principle, we quantify a population coding efficiency by fitting an exponential function to the error probability as a function of population size:

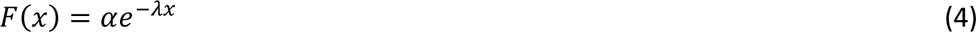

Where λ is the efficiency rate value and *x* is the neural population size. A higher efficiency rate value, the more efficient the population is at discriminating between conspecific stimuli presented from different spatial positions or orientations.

### 2D Activation Heatmaps

Two-dimensional activation heatmaps are valuable as a qualitative tool for visualizing differences in neural responses when the stimulus is presented from different spatial locations and orientations around the fish. We used a 3D model of a fish on which a population of electroreceptor locations have been placed (i.e., each dot on the fish) that was used previously (Ramachandra et al., 2023). The receptive field center of each neuron was delineated experimentally by moving a local dipole across the body surface and determining the edge of the classical receptive field where the neurons responded with their characteristic ON-center or OFF-center responses. All tuberous receptor locations within a neuron’s receptive field boundary were assigned to that neuron, such that one receptor could belong to several neurons’ receptive field centers. Values from the neural response measures (e.g., firing rate, gain, vector strength) for a given stimulus were appended to all receptors within the neurons’ receptive field centers, and then averaged so that each receptor’s location represented a single activation value for all its represented neurons.

## Results

Conspecific stimuli played from an artificial, conspecific dipole mimic (i.e., “fishpole”), were presented to an immobilized *A. leptorhynchus* while recording extracellularly from ELL pyramidal cells *in vivo*. After mapping the cell’s receptive field, the fishpole stimulus was positioned at one of three orientations (0,45,90°), and one of seven spatial positions (Fig 1A; see Methods) around the immobilized fish in random combination until all combinations of spatial stimuli were used. To replicate the signals experienced when a conspecific is present, we replicated the beat AM (i.e., sinusoidal amplitude modulations of the fish’s own EOD) and used, in this paper, a beat frequency of 30 Hz because all pyramidal cells respond strongly at this frequency.

We examined how the location or orientation of the stimulus fish would influence the pyramidal cell responses. We noticed obvious qualitative differences in the neural response that ranged across the spectrum of temporal to rate aspects of the spike train. The stimulus-elicited changes in response pattern were heterogeneous with some neurons showing pronounced changes in their mean firing rate whereas other cells responded with clear differences in vector strength with little to no change in rate (Fig 1B). We could not readily identify a single aspect of the response that most clearly correlated with changes in stimulus position. We therefore used three measures in our analysis that cover the range of rate vs temporal coding: mean firing rate, gain (which reflect changes in both timing and rate of the response), and vector strength (i.e., how tightly the response is concentrated at one phase of the stimulus).

Our aim is to compare how accurately the population of pyramidal cells encode that spatial location of the stimulus. To do so, we compare pair-wise population responses to different stimuli locations and quantify how different the pattern of responses are. The similarity in response pattern is based on a weighted Euclidean distance analysis that tightly correlates with the amount of information that the population carries about stimuli differences (i.e., location). This analysis results in an estimate of the error rate in stimuli discrimination that would occur by comparing population responses. This error rate is estimated as a function of the number of neurons included in the population response, a faster decrease in error rate with increasing population size (efficiency *λ* in Figure 2) indicates that the neurons carry more spatial information. We found that all unique pairs of spatial stimuli were able to be discriminated effectively, with the average of all unique stimulus pairs being discriminated reliably (<5% error) with populations of less than 20 neurons (Fig 2B). Thus, this data showed that the spatial aspect of the conspecific stimulus was reliably encoded within the pattern of the population response. Some neurons clearly changed their response pattern for different stimuli positions whereas others showed more subtle changes. By comparing subpopulations of neurons with clear difference for a given stimulus pair, to a subpopulation with less obvious differences, we demonstrate in Figure 2C how our efficiency measure reflects how accurately stimulus position is encoded in the response pattern.

**Figure 2.**
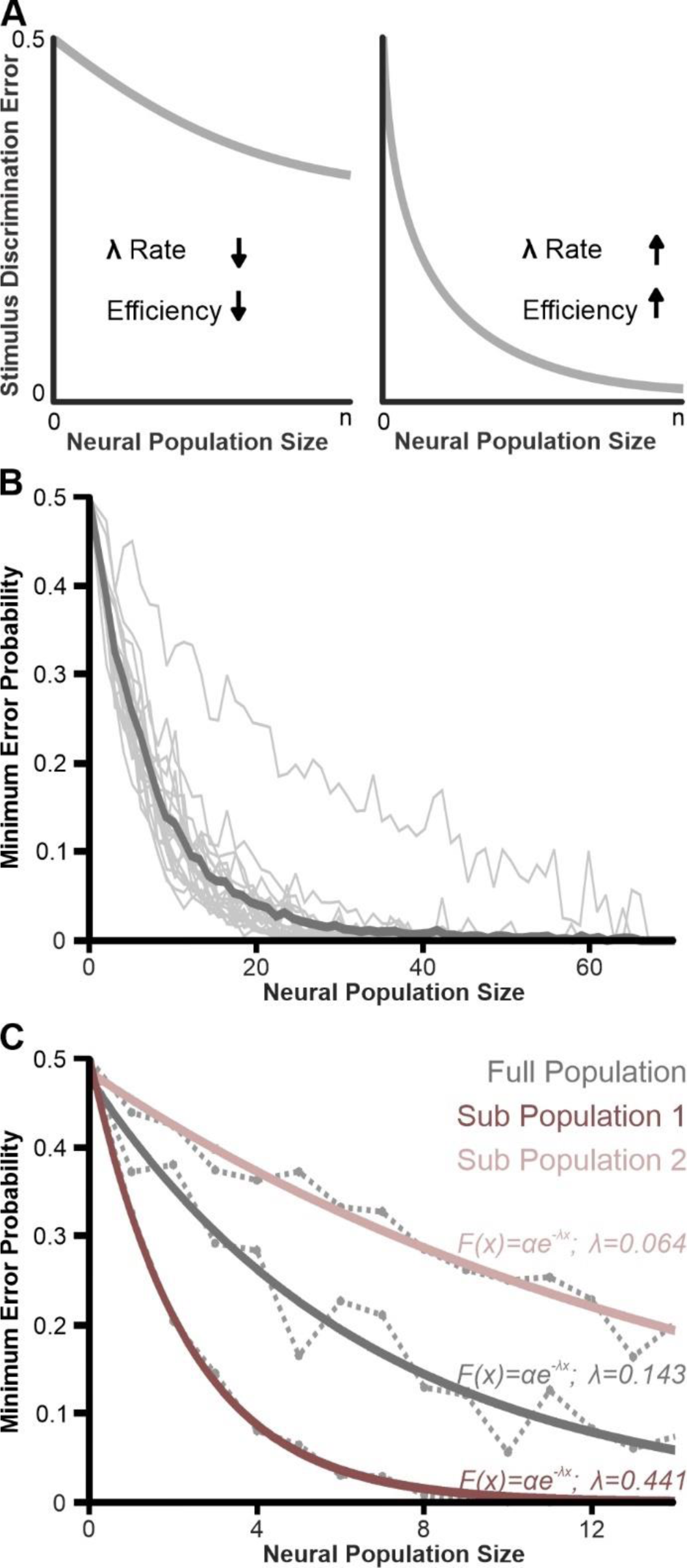
Coding efficiency quantified by the rate of decrease of the discrimination error as the information from more neurons are included in the analysis. **(A)** Schematic detailing how coding efficiency is related to the rate of discrimination error. For a pairwise stimulus discrimination task, the efficiency (λ) can be defined as the change in discrimination error as a function of neural population size. A slow decrease in error as the information from more neurons is pooled indicates a low coding efficiency (each neuron has little information or redundant information). A faster decrease (left plot compared to the right) indicates a higher efficiency. **(B)** Pairwise stimulus discrimination using vector strength on the full population of recorded pyramidal cells (n=70). All unique paired combinations of spatial stimuli are shown in the background (light gray), with the mean across all stimulus pairs in the foreground (dark gray). A discrimination accuracy level of 95% is obtained with fewer than 20 neurons. **(C)** Stimulus discrimination and efficiency across different neural populations. We selected two subsets of neuron (n=14 each): one where we could see obvious differences in responses between two stimuli locations and one where differences were not obvious. We used our analysis on these two subsets simply to illustrate the results expected from efficiently coding neurons (Sub-Population 1) and from a population with low coding efficiency.

ELL pyramidal cells are heterogenous and many differences in their response properties have been documented. Yet, it is not known whether the different subpopulations differ in their encoding of the spatial aspect of conspecific signals. To answer this question, we compare the coding accuracy across categories: ON-type vs OFF-type cells, neurons of the LS vs CLS maps, and superficial/intermediate vs deep pyramidal cells. ON and OFF-type pyramidal cells were easily distinguished based on their preference of stimulus polarity (increases vs decreases in stimulus amplitude). Location of each recorded neuron relative to the different ELL maps was estimated based on the stereotaxic position of the recording electrodes and on the response properties of the neurons (see Supplementary Figure S1). In this study, we focus on the LS and CLS segments that are most relevant for processing conspecific signals. Important differences exist between deep pyramidal cells and cells that are more superficial. We pooled together putative superficial and intermediate cells because they occupy a similar place in the circuitry of the electrosensory system whereas deep pyramidal cells are functionally separate (Maler, 2009b). We categorized deep-type pyramidal cells based on their characteristically high spontaneous firing rate, lower coefficient of variation, and their better synchronization to the EOD compared to superficial and intermediate-type pyramidal cells (S 1D; S 1E; Bastian, 1986; Bastian et al., 2002; Krahe et al., 2008).

The classical receptive field of each neuron was delineated based on their response to a small local dipole that was moved across rostro-caudal and ventro-dorsal locations. The neurons we recorded had receptive fields in various positions from head to tail (Fig 3A, 3B). We note that, while most electrophysiological studies avoid sampling cells from the fish’s head because the typical experimental configuration has the fish’s head close to -or above-the water surface, we performed the experiment with the fish completely submerged in a more realistic position. We did not observe striking differences in response properties for cells of the head, despite the fact that they receive inputs from much more densely pack receptors than pyramidal cells from the trunk of the fish. Receptive field size varied from cell to cell (Fig 3B). As expected, CLS neurons had smaller receptive field sizes than LS neurons (Carr et al., 1982; see also Supplementary Figure S2). The receptive field sizes for deep-type and superficial/intermediate-type pyramidal cells were similar (Maler, 2007; Maler, 2009b; Maler, 2009a).

**Figure 3.**
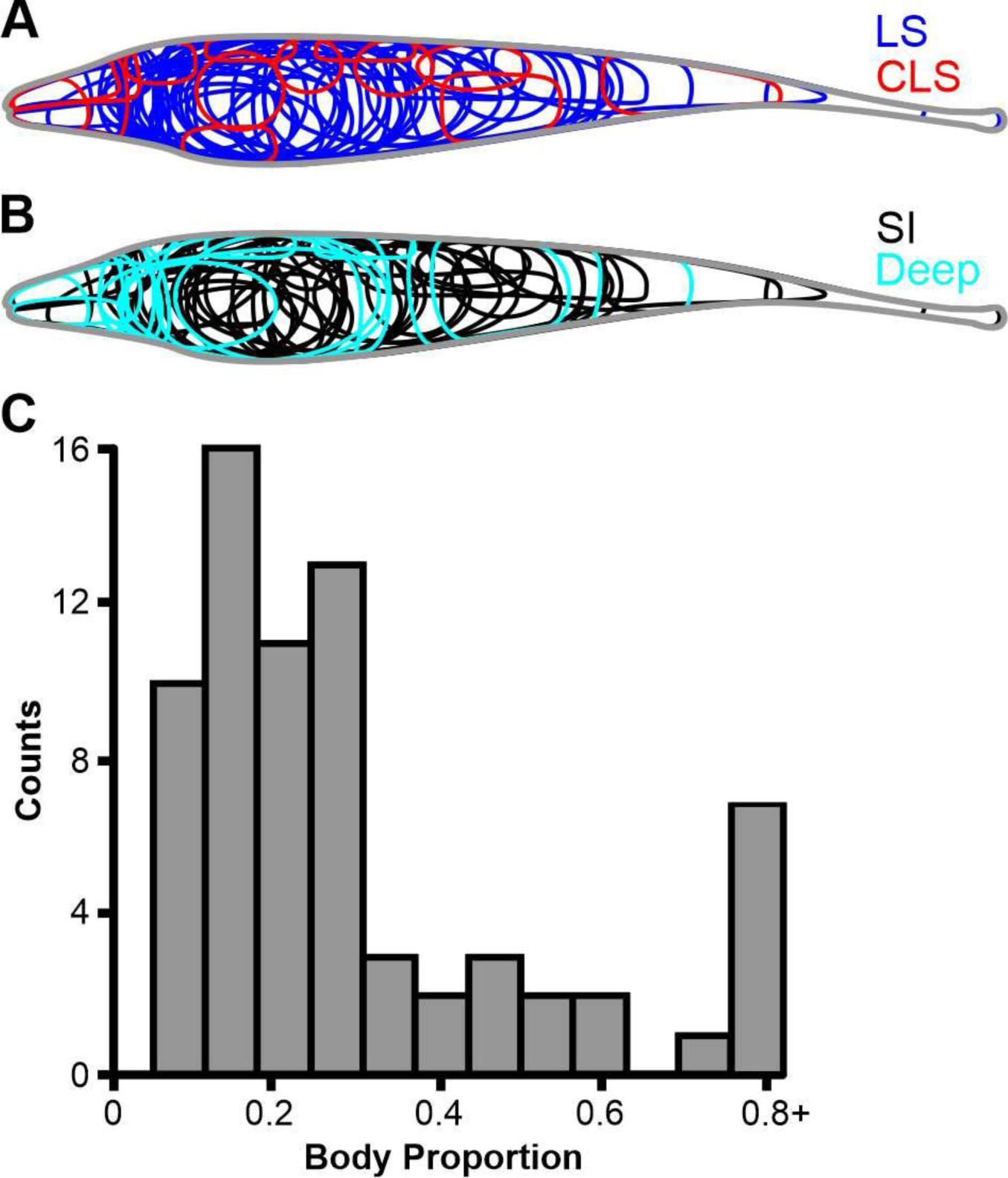
Receptive field of pyramidal cells sampled. **(A)** Boundaries of receptive field centers from recorded LS (n=55) and CLS (n=15) pyramidal cells on a two-dimensional outline of an *A. leptorhynchus*. **(B)** Boundaries of receptive field centers from recorded deep-type (n=18) and superficial/intermediate-type (n=52) pyramidal cells. **(C)** Histogram of all recorded pyramidal cell receptive field sizes (n=70). Size is represented as a fraction of the length in the rostro-caudal axis compared to the total body length.

To visualize how ELL pyramidal cells varied in their response to stimuli from different locations, we constructed two-dimensional activation heatmaps based on one of the three response strength measurements (see Fig 1). These heatmaps (Fig 4) highlight a few key observations. First, as expected, stimuli from different locations lead to clear differences in the pattern of activation across the body. Also, the heatmaps highlight the heterogeneity and variability in the response pattern of pyramidal cells. Although an overall pattern of activation is visible, with receptive field facing the stimulus location being more strongly activated, pyramidal cells showed uneven patterns of activation with movement relative to their receptive field. Finally, these heatmaps illustrate that different subpopulations of pyramidal cells might have a more obvious relationship between their response strength and the stimulus location. Specifically, we displayed responses of ON vs. Off and deep vs. superficial/intermediate where the ON and the deep cells show clearer differences across stimuli locations.

**Figure 4.**
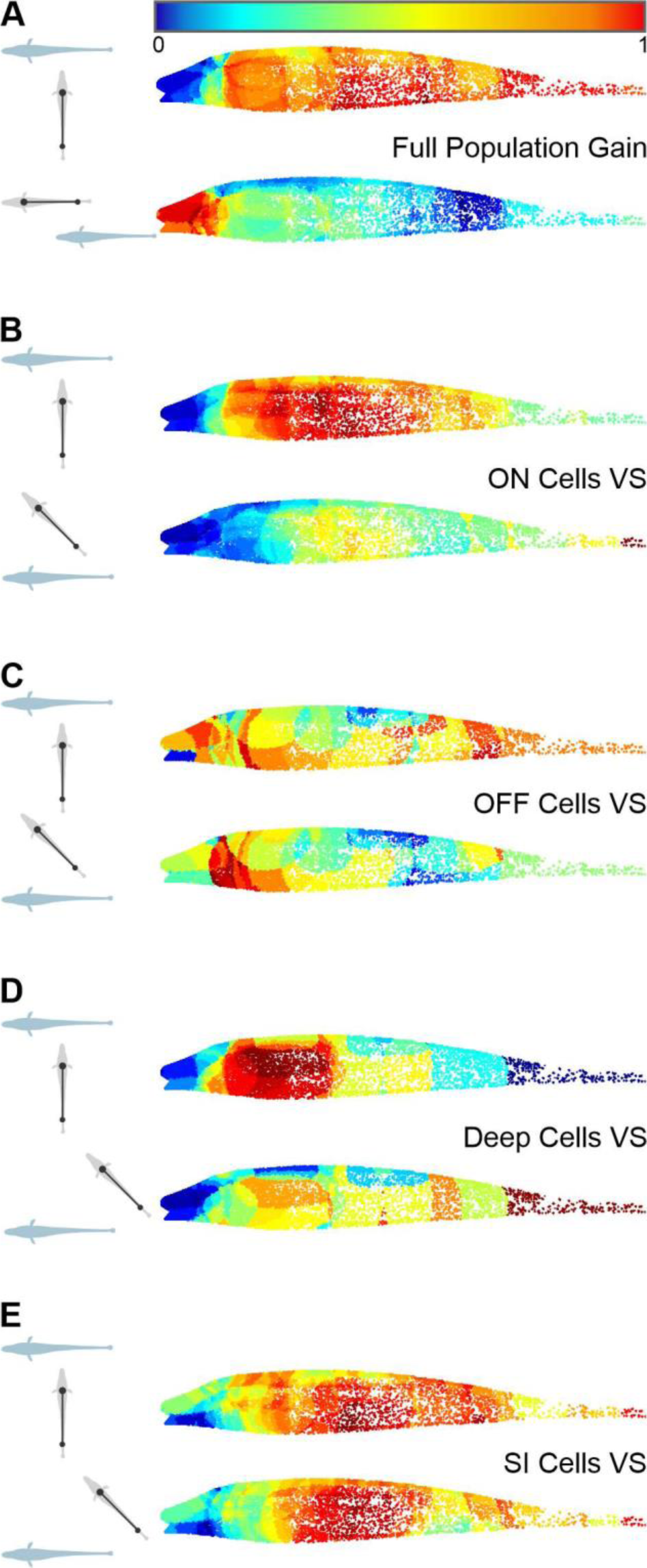
Responses to spatially realistic conspecific signals visualized as topographic heatmaps for different subsets of pyramidal cells. The heatmaps allow a visualization of the population response to a conspecific stimulus played from various relative positions and orientations (shown on the left insets). Each colored point on the heatmap represents a putative receptor on the skin of the fish (see Methods and Ramachandra et al., 2023). A receptor can contribute to several neurons’ receptive field and its color will reflect the average responses (e.g., gain) across these neurons. For each neuron to contribute equally to the heatmap, their responses are normalized to 1 where 1 is the strongest response of the neuron across all stimuli positions. In the 5 pairs of heatmaps presented here for different subpopulations of cells, we see differences across stimulus locations that vary from obvious (A, B, D) to more subtle (C, E) **(A)** Heatmaps of the full population of recorded pyramidal cells (n=70) using gain as a response measure. **(B)** Heatmaps of the ON-type pyramidal cells subpopulation (n=46) using vector strength as response measure. **(C)** Heatmaps of the OFF-type pyramidal cells subpopulation (n=24) using vector strength as response measure. **(D)** Heatmaps of the deep pyramidal cells subpopulation (n=18) using vector strength as response measure. **(E)** Heatmaps of the superficial/intermediate pyramidal cells subpopulation (n=52) using vector strength as response measure.

Further analysis revealed that the discrimination error for specific spatial stimulus pairs varied based on the response measure used in the analysis. Overall, we found that using the mean firing rate of the vector strength to quantify the response led to better spatial coding than using the gain (Fig 5A). It is worth mentioning that we also investigated how the phase of the response changed with the spatial stimulus. Some neurons showed noticeable shifts in response phase to a conspecific stimulus changing either orientation or location (Supplementary Figure S3). However, this measure proved less informative for population analysis, when their responses were combined with either cells that also exhibited phase changes or cells that showed no phase changes in their response.

**Figure 5.**
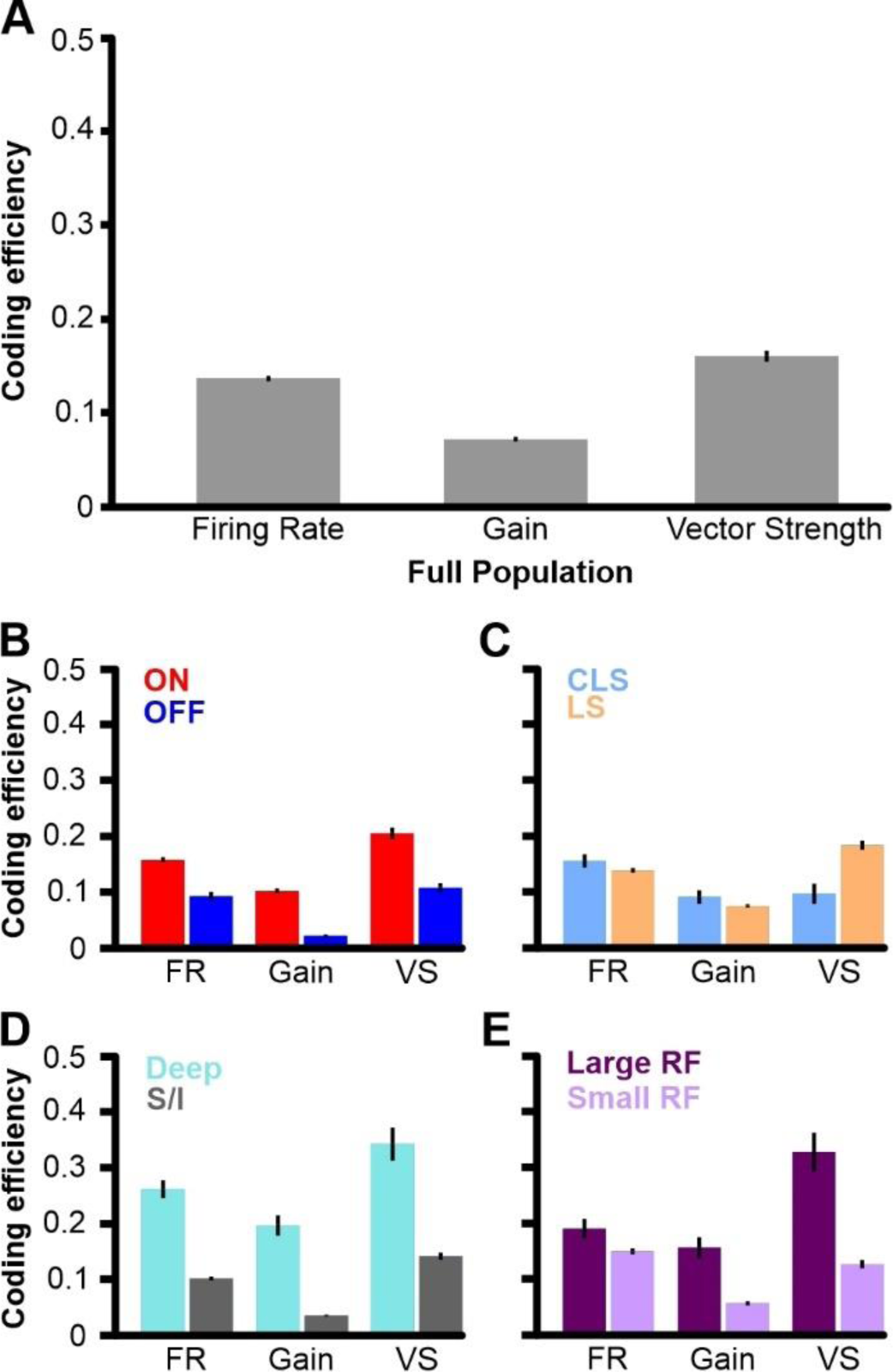
Spatial coding efficiency varies with pyramidal cell type and is dependent on the aspect of the neural response relevant for stimulus discrimination. **(A)** Mean coding efficiency (± s.e. across stimuli pairs) of all pairwise stimulus combinations (orthogonal orientation; see Methods) using the full population (n=70). The highest efficiency is obtained when using vector strength to characterize neural responses; and the lowest efficiency is obtained when using response gain (*p* < 0.0001). **(B)** Mean coding efficiency (± s.e. across stimuli pairs) for populations of ON (n=46) and OFF-type pyramidal cells (n=24). The highest efficiency results from vector strength, with the lowest efficiency from stimulus-response gain (ON - *p* < 0.0001, OFF - *p* < 0.0001). ON-type pyramidal cells have higher efficiency than OFF-type pyramidal cells across all measures (*p* < 0.0001). **(C)** Mean coding efficiency (± s.e. across stimuli pairs) for populations of CLS (n=15) and LS pyramidal cells (n=55). There is a higher efficiency result from mean firing rate compared to gain for CLS (*p* < 0.0001), and for vector strength compared to gain for LS (*p* < 0.0001). **(D)** Mean coding efficiency (± s.e. across stimuli pairs) for populations of deep (n=18) and superficial/intermediate-type pyramidal cells (n=52). The highest efficiency results from vector strength, with the lowest efficiency from stimulus-response gain (Deep - *p* < 0.0001, SI - *p* < 0.0001). Deep-type pyramidal cells have higher efficiency than superficial and intermediate-type pyramidal cells across all measures (*p* < 0.0001). **(E)** Mean coding efficiency (± s.e. across stimuli pairs) for cells with large receptive fields (n=33) and small receptive fields (n=37). The highest efficiency results from using vector strength for large receptive field neurons (*p* < 0.01), and is highest using mean firing rate for small receptive field neurons (*p* < 0.01). Large receptive field pyramidal cells have higher efficiency than small receptive field pyramidal cells across all measures (*p* < 0.0001).

The full population was separated into distinct categories of pyramidal cell types, and we asked if specific pyramidal cell types would discriminate spatial stimuli more efficiently depending on the aspect of the spike train response used for discrimination. We found that ON-type pyramidal cells can discriminate the spatial stimulus more efficiently than OFF-type pyramidal cells across all three measures (Fig 5B). Similar to the full population, both ON and OFF-type pyramidal cells obtained the highest efficiency rate using the vector strength, and the lowest when using stimulus-response gain. This exact finding was also observed when comparing deep-type pyramidal cells vs superficial/intermediate-type pyramidal cells (Fig 5D). However, when comparing efficiency between CLS and LS pyramidal cells, there was a flip in the measures that resulted in the highest efficiency rate. The population of CLS neurons was found to be most efficient when using mean firing rate, whereas the LS population was most efficient using vector strength (Fig 5C). The same trend is observed when comparing neurons with large receptive fields (> 0.2 body proportion) against neurons with small receptive fields (< 0.2 body proportion; Fig 5E). This might reflect the fact that LS neurons tend to have larger receptive fields than CLS neurons. Our data thus indicates that CLS and LS neurons encode spatial information with a different proportion of rate vs. temporal coding, with CLS cells relying more on rate coding compared to LS cells that encode spatial information better in the timing of the response.

An alternative approach to characterizing how different neurons encode spatial information is to describe the properties of neurons that carry relatively more information about stimulus location. Our decoding analysis assigns a weight to each neuron based on how different their response is to the stimuli locations being compared. This weight is based on the Kullback-Leibler divergence between response distributions which relates to the information present in the spike trains (Allen and Marsat, 2018; Allen and Marsat, 2019; Marsat and Maler, 2010; Marsat et al., 2023; van Rossum, 2001). By using the average weight assigned to a neuron across stimuli comparisons, we categorized the cells as being associated with high weights (> 0.7; Fig 6A) or low weights (< 0.7). Consequently, the cells in each group have a higher vs. lower coding efficiency (Fig. 6B). This approach is complementary to our previous analysis because it allows us to determine the characteristics of cells that encode spatial information particularly well. We found that the high-weight group carried more information in their response timing (i.e., vector strength; Fig 6B), whereas the low-weight group encoded more spatial information in their mean firing rate. Interestingly, this trend parallels the differences we observed comparing LS and CLS cells and cells with large vs. small receptive fields (see Fig 5). Our analysis of cell properties in each weight category also confirms the previous findings. Specifically, cells in the high weight category had a higher average spontaneous firing rate, lower average coefficient of variation, and larger average receptive field size than neurons in the low weight category (Fig 6C; Fig 6D; Fig 6E). We verified that putative deep pyramidal cells were more likely to be in the high-weight category (Fig 6F).

**Figure 6.**
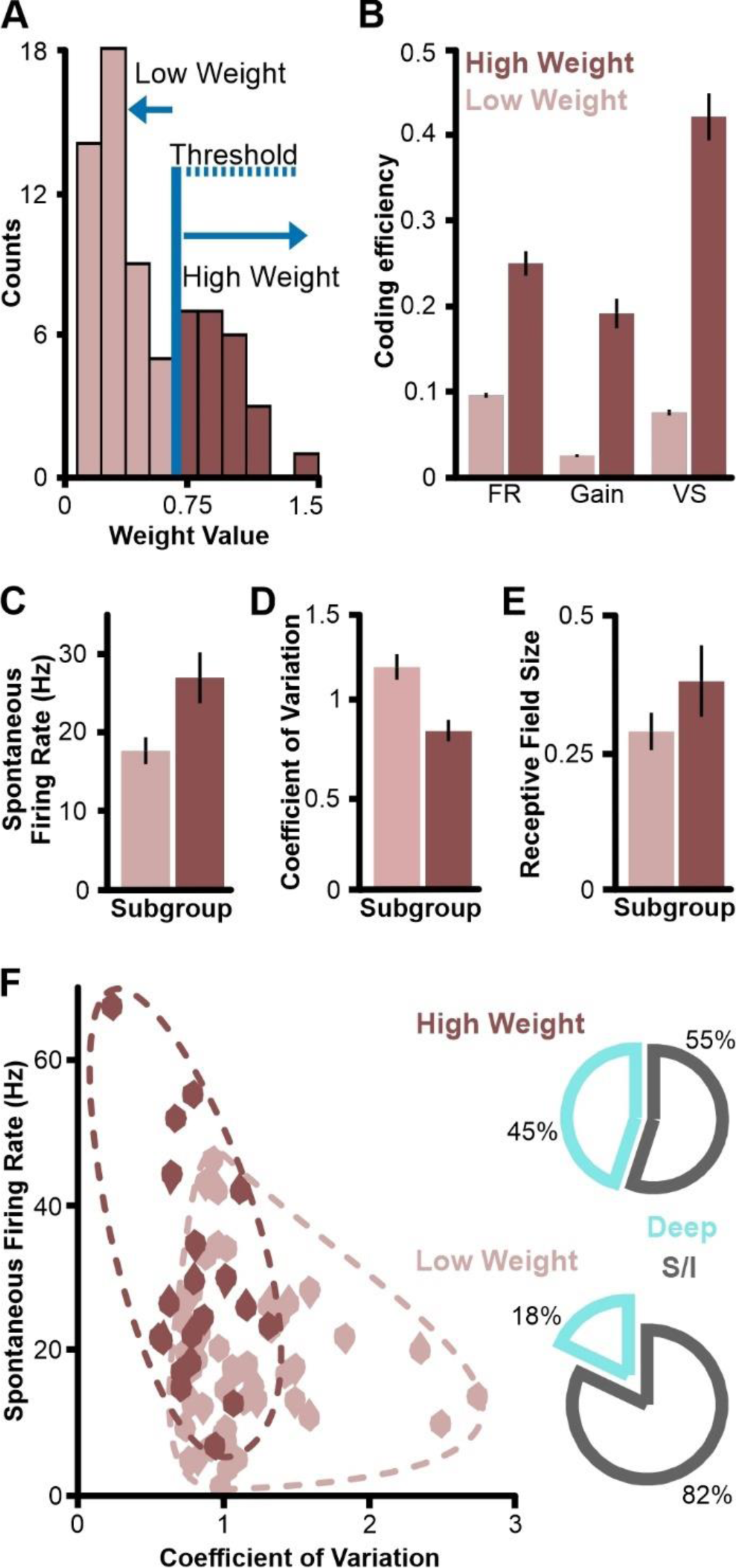
Properties of neurons with more informative responses. **(A)** Distribution of average weight value assigned to each neuron in the analysis that reflects the separation in their response distribution to the stimuli being compared (here averaged across stimuli pairs). A threshold was established to divide this distribution with two peaks into two different populations of neurons (low weight in pink, n=50; high weight in red, n=20). **(B)** Mean coding efficiency (± s.e. across stimuli pairs; orthogonal orientations only), of the two weight-groups show the expected overall difference, the high-weight performs much better (*p* < 0.01). **(C)** The mean spontaneous firing rate (± s.e. across neurons) for the high-weight group is higher than that of the low-weight neurons (*p* < 0.01). **(D)** The mean coefficient of variation (± s.e. across neurons) for the high-weight group is lower than that of the low-weight neurons (*p* < 0.001). **(E)** The average receptive field size (± s.e. across neurons) for the high-weight group is higher than that of the low-weight neurons, though this difference is not statistically significant (*p* ∼< 0.05). **(F)** Scatter plot of spontaneous firing rate and coefficient of variation of every neuron in each weight category. We delineated the groups with a dashed line to highlight the separation/overlap between groups. Pie charts showing the proportion of deep pyramidal cell types within each weight category (right). Note that only 18 of 70 recorded neurons are deep pyramidal cells which represents 25.7%. deep pyramidal cells are thus under-represented in the low-weight group and over-represented in the high-weight group.

The results presented in previous figures averaged the analysis of pairs of stimuli locations. We now ask if the difficulty in discriminating between spatial stimuli could be influenced by the proximity of the stimulation sites. For example, two spatial stimuli that are close to one another may be a more difficult discrimination task than two spatial stimuli that are far apart. We hypothesized that the differences we observed in spatial coding efficiency across cell types were even more apparent when considering only the more difficult discrimination tasks. Surprisingly, we found only modest differences in coding efficiency when comparing stimuli that are next to each other (ipsilateral, Fig 7) or on opposite sides of the body (contralateral). Overall, coding efficiency was better for coarse discrimination (contralateral) than fine discrimination (ipsilateral). The cell sub-type performing best, and the response properties encoding the most information were the same as noted in the previous analysis and there were no qualitative differences when considering coarse vs. fine discrimination tasks. Our stimuli locations and intensity mimicked a medium-sized fish relatively close to the fish being tested. Recent finding suggests that the spatial information present in the population of receptors should be relatively accurate for stimuli at these distances (Ramachandra et al. 2023). It is thus probable that even when comparing our closest stimuli locations, we are not testing the lower limits of the cell’s sensitivity. Using weaker stimuli (Supplementary Figure S5), placing the stimulus source further away or comparing locations with less separation would result in lower coding performance and potentially increase the modest differences we observe between fine and coarse discrimination tasks.

**Figure 7.**
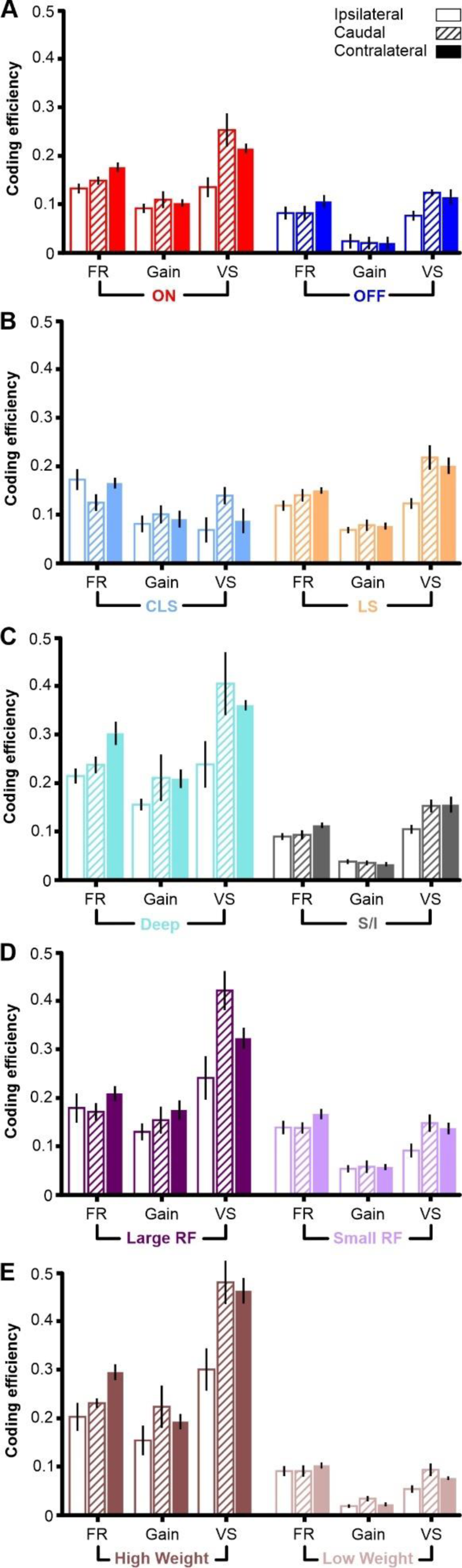
Fine and coarse spatial discrimination across cells-types and response measures. Mean coding efficiency (± s.e. across stimuli pairs) for discrimination tasks where we compare: locations on the same side of the fish (ipsilateral); locations on opposite sides (contralateral); or locations on the sides compared to the caudal location. We compare different subpopulations and groups of pyramidal cells: **(A)** ON-type vs. OFF-type pyramidal cells; **(B)** CLS vs. LS, **(C)** deep-type vs. superficial/intermediate-type pyramidal cells; **(D)** large receptive field vs. small receptive field neurons, **(E)** high weight vs. low weight neurons.

## Discussion

We investigated spatial coding of conspecific stimuli by electrosensory neuron populations in the hindbrain. To do so, we performed *in-vivo* electrophysiological recordings on immobilized *A. leptorhynchus*, as an artificial dipole mimic (i.e., “fishpole”) was used to stimulate from various positions within the experimental tank. Specifically, we targeted the topographic maps of the hindbrain ELL, which contain heterogeneous populations of pyramidal cells that vary in their anatomy and physiological response properties. Using our full dataset of recorded neurons, we found that this full population could discriminate accurately (< 5% error) between all unique pairs of spatial stimuli presented (see Methods). In addition to heterogeneity in anatomy and physiology, ELL pyramidal cells also vary in their functional connectivity based on their layer in the map. For example, pyramidal cells in the deep layer have no receptive field surround and do not receive feedback from either the nucleus praeminentialis or via cerebellar granular cells from the posterior eminentia granularis. In fact, deep pyramidal cells are the source of feedback to superficial and intermediate-type pyramidal cells in the ELL (Maler et al., 1991; Berman and Maler, 1999; see also Milam et al., 2019, for review). We therefore investigated how the discrimination ability differs between subpopulations in the ELL. Briefly, we found that: (1) ON-type pyramidal cells displayed lower error than OFF-type; (2) deep pyramidal cells outperformed superficial and intermediate-type; (3) pyramidal cells with larger receptive fields were better than those with small receptive fields; and (4) LS neurons discriminated more accurately than CLS neurons on average. By far, the clearest difference we observed in spatial coding was between the deep and superficial pyramidal cell populations. Previous studies have shown that superficial pyramidal cells excel in temporal coding of communication signals (e.g., chirps). Interestingly, in this study we found that superficial cells performed poorly in encoding spatial information from conspecific signals. When contrasted with deep cells, which excelled in the same spatial coding task, this result complements findings from Vonderschen and Chacron (2011). In their study, they describe a dichotomy of sparse and dense coding strategies by neural subpopulations, downstream from the ELL in the midbrain torus semicircularis. Sparse coders in the torus were specialized for encoding sensory information related to specific chirp features, whereas dense coders were more broadly responsive to electrosensory stimuli. Similar to the torus, the ELL has a laminar structure, as well as a complex network of connections. Electrosensory input from the ELL to the torus is topographically conserved and confined to the dorsal torus (Carr et al., 1981). Downstream, the electrosensory pathway separates as the torus outputs: to the optic tectum involved in spatial processing; to the nucleus electrosensorius involved in processing communication signals; and to the preglomerular nucleus that mediates connectivity with the forebrain (Giassi et al., 2012; Zupanc and Horschke, 1997a; Zupanc and Horschke, 1997b). Topography is conserved to brain regions as far as the optic tectum, but is lost in the nucleus electrosensorius, the preglomerular nucleus, and the forebrain dorsal telencephalon (pallium). It is for this reason that the optic tectum is considered to be an important area for multisensory integration, and putatively an ultimate localization center. Taken together, our findings suggest that the separation of spatial vs identity (e.g., communication) coding begins as early as the ELL with an early split starting between deep vs superficial pyramidal cells, albeit a large overlap in function. Further studies are needed to investigate spatial coding in higher brain areas where topography remains conserved.

Our results show that certain neural subpopulations allow for more accurate discrimination when using mean firing rate, while others encode more accurately in their response synchrony. We found that this pattern is most obvious when comparing population responses between the LS and CLS maps. Pyramidal cells in the LS map displayed less discrimination error when using vector strength, whereas CLS cells performed better, on average, when using mean firing rate. This response preference is indicative of a switch in the neural coding strategies used by different topographic maps. Several factors might be contributing to the differences in coding preference that we observed here, such as: receptive field parameters, adaptation, and the influence of feedback. Nonetheless, this finding provides an opportunity for speculation as to what measures of the spiking response are most relevant for spatial coding in downstream sensory areas. For example, it is unclear what aspects of the pyramidal cells’ response influence spatial coding in the midbrain torus. While we know that dense coders in torus will modulate their firing rate with peak and trough of a conspecific beat, the strength of the response can be affected by the average number of spikes or by having a spike that clusters more around a single phase, such that both spike rate and timing could be relevant. Preferences in the neural response tailored to specific stimulus features have been well documented, some good examples being combinatorial and multiplexed neural codes (Bodnar and Bass, 1999; Lankarany et al., 2019). For example, studies on human sound localization have shown how neurons that receive shared input can use asynchronous firing rate to encode the intensity of low-contrast features, while also using precise timing of synchronous spikes to encode high-contrast features (Lankarany et al., 2019). Similarly, other behavioral experiments on human sound localization have found that softer sounds can be perceived closer to the midline than louder sounds, favoring a rate-coding strategy (Ihlefeld et al., 2019). Furthermore, research on spatial navigation has shown that the time of firing can represent an animal’s location within a place field, whereas the firing rate can represent the animal’s velocity through the field (Huxter et al., 2003). Information can also be transmitted through short interspike intervals within a burst (Krahe and Gabbiani, 2004; Oswald et al., 2007). Thus, it is well supported that in heterogeneous neural populations spatial information about a conspecific’s location can be represented in different aspects of the spiking response. Indeed, further studies are necessary to dissect the role of information coding related to conspecific location.

Our results demonstrate that spatial coding efficiency is high across most subpopulations of pyramidal cells in the ELL. Accurate discrimination between pairs of stimulus positions is possible using a small number of cells relative to the full population. Specifically, the discrimination accuracy is high for conspecifics located at distances of 10 cm away. At this distance, we are not testing stimuli at the edge of sensitivity, as recent studies have estimated that difficult discrimination tasks start at distances of approximately 30 cm away (Ramachandra et al., 2023). Furthermore, our discrimination analysis takes 1 s averages of neural responses and assumes that a decoder can integrate these inputs over time. If integrating over less time, because the neural system does not operate on long time scales or because the fish does not remain in a fixed location, the discrimination accuracy will decrease. Moreover, this decoding analysis might not encompass the relevant aspect of the neural response, or could underestimate coding accuracy. An alternative decoding method might implement a principal component approach to average out noise more effectively. On the other hand, our analysis could also be overestimating the coding efficiency. In our analysis, we weigh each neuron and thus leverage the most informative neurons over those that provide less information. It is possible that this may not be the exact computation that subpopulations of ELL pyramidal cells are performing, as our measure relates directly to the amount of information present in the system (Marsat et al., 2023). Additionally, the gathering of information can be enhanced via active sensing behaviors. Such specialized and often stereotype-patterned behaviors occur across systems and include edge detection and tracking of odor plumes in moth olfaction, and foveal sampling in the visual and electrosensory systems, to name a few (Enikolopov et al., 2018; Pedraja et al., 2019). Certain bat species have been shown to take advantage of their angle of approach with respect to the background surface to increase the signal to noise ratio of a prey echo during prey capture behavior. Such acoustically camouflaged prey items would normally have their weak prey echoes masked by background echoes from other objects in the natural environment (Geipel et al., 2019). High accuracy steering towards the location of a sound source at a fixed azimuth has been documented in crickets (Schöneich and Hedwig, 2010). This finding suggests that localization ability could be high when integration over time can happen. Cricket zig-zag walking and other corrective repositioning behaviors, could help to re-evaluate the localization error during movement. Thus, localization during behavior should consider the motion component and that the process from sensory to motor and back is highly dynamic. Further studies on the role of active sampling and dynamic sensorimotor adjustments are needed to better understand how the spatial aspect of signals are encoded by the nervous system.

Our results provide new insights for population coding of spatially realistic conspecific signals and what aspects of the neural response are most important for localization. Overall, our data suggests that the start of segregation of spatial processing occurs in ELL pyramidal cells. It is likely that the experimental findings we present here for quantifying spatial coding performance in pyramidal cells are generalizable to other sensory systems. The neural circuitry in the ELL contains several network elements that are shared across modalities, such as classical receptive field center-surround organization, topographic maps of the body, and feedback influences that contribute to shaping the neural code. Taken together, this study serves as foundational work for understanding how a primary sensory area of the hindbrain represents the location of conspecifics. Future studies will be necessary to gain a better understanding of the complex interaction between extracting the location and the “message” in sensory signals.

## Supplementary Information

**Supplemental Figure 1.**
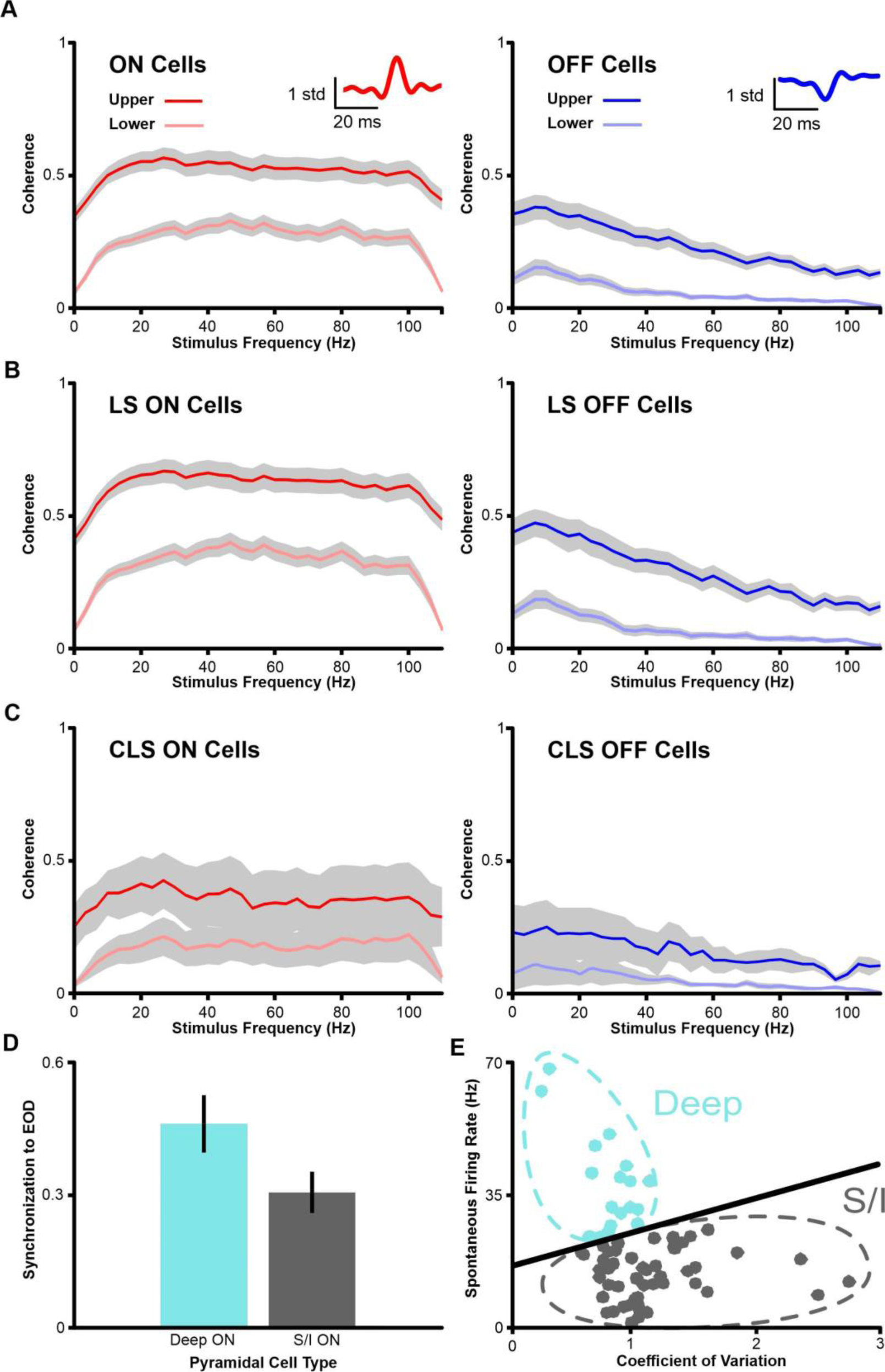
Confirmation of pyramidal cell type. **(A)** Upper and lower bound coherence of ON and OFF-type pyramidal cells. Insets show spike triggered average waveforms in response to RAM stimuli presented globally. Coherence analyses are standard and described in previous publications (Allen et al., 2019; Krahe et al., 2008). The upper-bound coherence reflects the coding accuracy including both linearly and non-linearly encoded information, whereas lower-bound coherence is based on the linear correlation between the stimulus and the response, gray shaded areas represent ±1 s.d. across neurons. **(B)** Upper and lower bound coherence of LS ON and LS OFF-type pyramidal cells. **(C)** Upper and lower bound coherence of CLS ON and CLS OFF-type pyramidal cells. **(D)** Synchronization to the EOD between deep and superficial/intermediate-type pyramidal cells. The synchronization uses the vector strength measure (ranging from 0 to 1) in response to cycles of the EOD rather than cycles of a SAM stimulus. Deep-type pyramidal cells show higher EOD phase locking (*p <* 0.05). Vertical, black lines indicate ±1 s.e. **(E)** Scatterplot of the baseline firing rate and coefficient of variation for each neuron recorded from the full population (n=70). Deep-type pyramidal cells are shown in blue, outlined manually for visualized grouping. Superficial and intermediate-type pyramidal cells are shown in black with manually outlined grouping to better visualize clustering of cells.

**Supplemental Figure 2.**
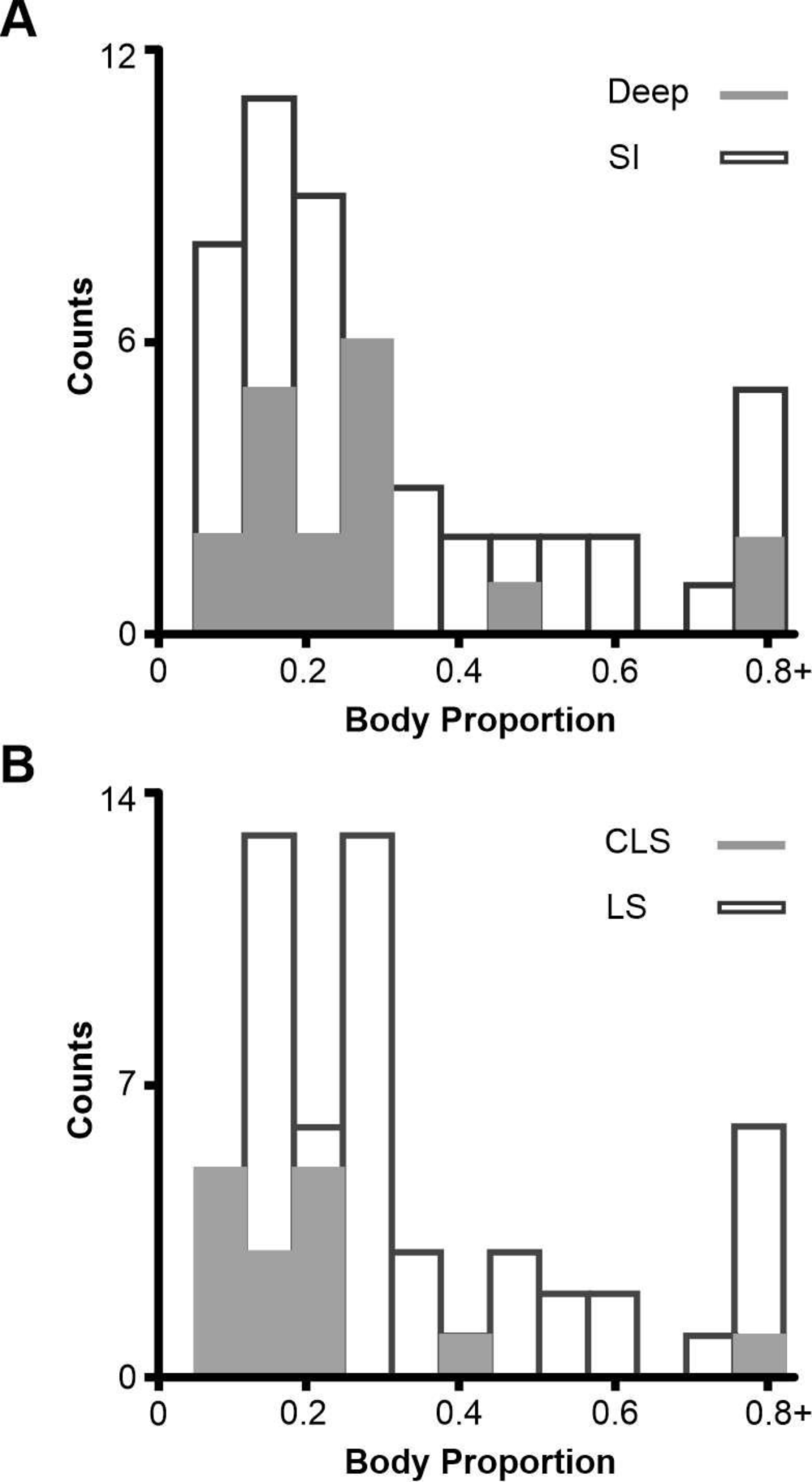
Receptive field size by pyramidal cell type. **(A)** Histogram of receptive field sizes measured as a fraction of total body proportion for deep and superficial/intermediate-type pyramidal cells. **(B)** Histogram of receptive field sizes measured as a fraction of total body proportion for LS and CLS pyramidal cells.

**Supplemental Figure 3.**
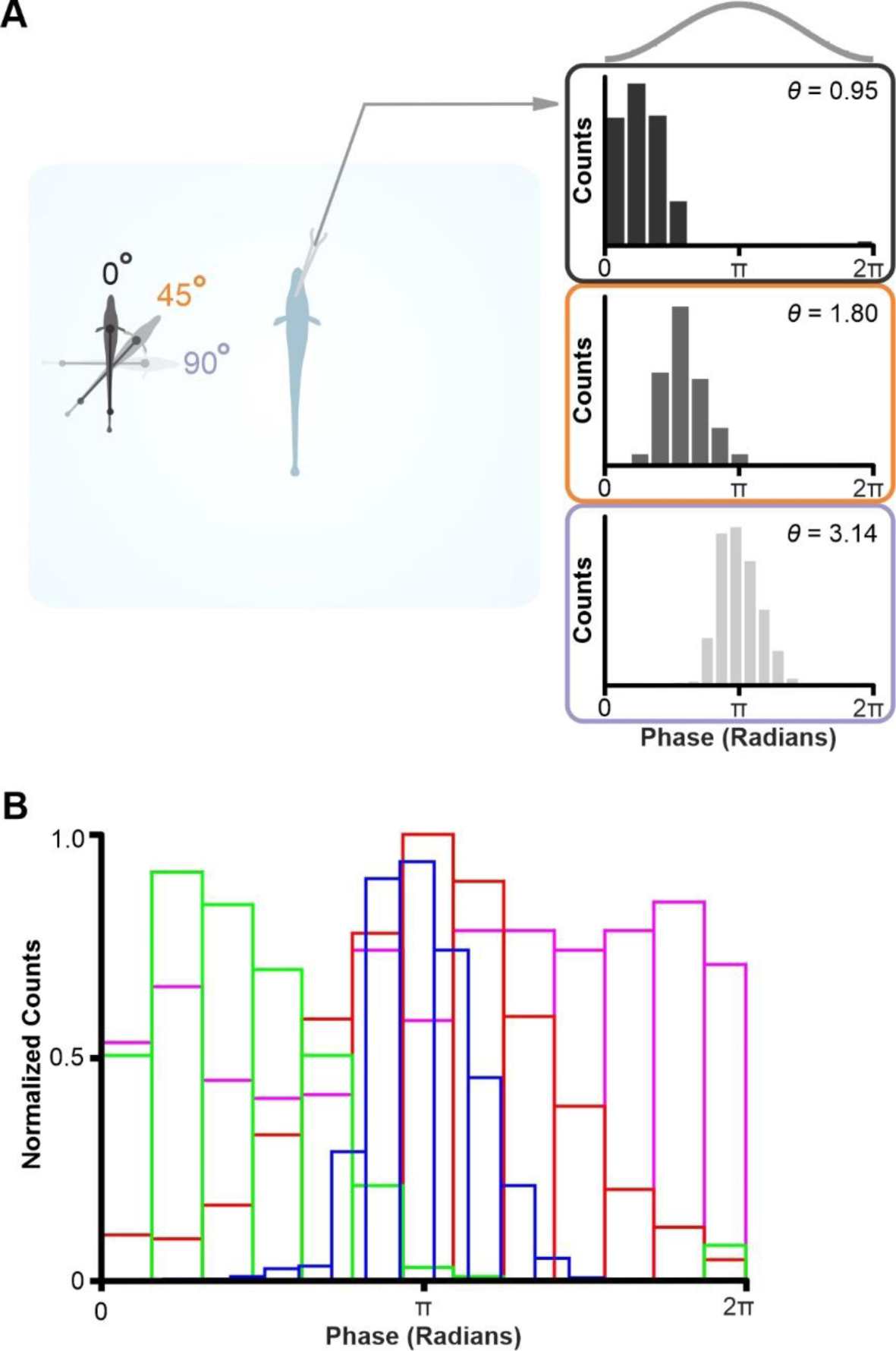
Observed phase-shifted responses in ELL pyramidal cells. **(A)** Effect of stimulus orientation a single ON-type ELL pyramidal cell. Certain neurons showed clear changes in phase to the spatially realistic conspecific stimulus. The average phase in the response is represented as θ. **(B)** 4 ON-type ELL pyramidal cells and their phase response to a stimulus with orthogonal orientation and placed in a singular location ipsilaterally to the receptive field. Each pyramidal cell response is shown as an unfilled histogram in color scale (in similar fashion to A).

**Supplemental Figure 5.**
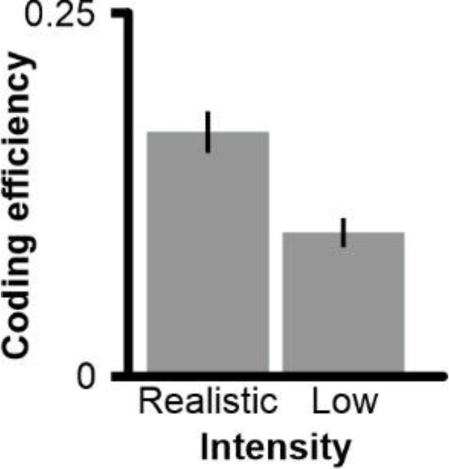
Fishpole intensity and its effects on discrimination efficiency. Discrimination efficiency of pyramidal cells (n=23) to the spatial stimulus using two different stimulus intensities. Discrimination occurs even a

## Notes

Funding: This research was supported by the NSF’s CAREER grant IOS-1942960 to GM.

### Competing Interest Statement

The authors have declared no competing interest.

